# Frequent development of broadly neutralizing antibodies in early life in a large cohort of HIV-infected children

**DOI:** 10.1101/2021.05.07.443081

**Authors:** Amanda Lucier, Youyi Fong, Shuk Hang Li, Maria Dennis, Joshua Eudailey, Ashley Nelson, Kevin Saunders, Coleen K. Cunningham, Elizabeth McFarland, Ross McKinney, M. Anthony Moody, Celia LaBranche, David Montefiori, Sallie R. Permar, Genevieve G. Fouda

**Affiliations:** Duke University Medical Center, Durham, NC; Statistical Center for HIV/AIDS Research and Prevention, Fred Hutchinson Cancer Research Center, Seattle, WA; University of California, Irvine; University of Colorado, Aurora, CO; Weill Cornell School of Medicine

## Abstract

Recent studies conducted in small cohorts of children have indicated that broadly neutralizing antibodies (bnAbs) may develop earlier after HIV infection compared to adults. To define the frequency and kinetics of bnAb responses in a larger pediatric cohort, we evaluated plasma from 212 ART-naïve, children living with HIV aged 1 to 3 years. Neutralization breadth and potency was assessed using a panel of 10 tier-2 viruses and compared to those of adults with chronic HIV. Further, the magnitude, epitope specificity and IgG subclass distribution of Env-specific antibodies were also assessed using a binding antibody multiplex assay. We found that 1-year-old children demonstrated neutralization breadth comparable to that of chronically-infected adults, and breadth continued to increase with age such that the pediatric cohort overall exhibited significantly greater neutralization breadth than adults (p= 0.014). Similarly, binding antibody responses increased with age, and the levels in 2 to 3 year-old children were comparable to those of adults. Overall, there was no significant difference in antibody specificities or IgG subclass distribution between the pediatric and adult cohorts. Interestingly, the neutralization activity was mapped to a single epitope (CD4 binding site, V2 or V3 glycans) in only 5 of 38 pediatric broadly neutralizing samples, suggesting a polyclonal neutralization response may develop in most children. These results contribute to a growing body of evidence suggesting that the early life immune system may present advantages for the development of an effective HIV vaccine.

**Author summary:** Understanding the development of broadly neutralizing antibodies during natural HIV infection is critical to provide insights for the development of an effective vaccine. In this study, we used a large cohort (n=212) of HIV infected children aged 1-3 years old to define the frequency and kinetics of broad neutralization in comparison to adults with chronic HIV. We also evaluated the magnitude and epitope specificity of HIV Env-specific antibodies in children and adults. We observed that both the binding and neutralizing antibody responses increased with age. However, by one year of age, children had neutralization breadth comparable to that of chronically infected adults; and overall, 1-3 year-old children demonstrated higher neutralization breadth than adults. Only slight differences in epitope-specificity and IgG subclass distribution were observed between children and adults, suggesting that these factors are not major contributors to the observed age-related difference in neutralization breadth. The epitope specificity of the neutralizing antibodies could not be mapped in the majority of children, suggesting that their response may be polyclonal. Our study contributes to the body of work indicating a potential advantage of the early life immune system for the elicitation of broad neutralization.

## INTRODUCTION

The global HIV pandemic persists despite increased availability of antiretroviral (ARV) treatment. In 2019, 1.5 million people aged 15 years and above became newly infected with HIV and 460,000 of these infections occurred in young adults aged 15-24 [1]. Furthermore, adolescents represent the only age group that has experienced increased numbers of AIDS-related deaths over the last decade [2]. While the need for an HIV vaccine is clear, it is critical that this vaccine be designed for administration prior to sexual debut to both protect during the vulnerable window of adolescence, as well as generate lifelong immunity.

Passive immunization studies in animal models have demonstrated that antibodies capable of neutralizing HIV-1 can protect from virus acquisition [3]. Elicitation of broadly neutralizing antibodies (bnAbs) is therefore believed to be crucial for vaccine efficacy. However, traditional vaccine approaches have failed to induce bnAb responses [4]. Furthermore, only a subset of chronically HIV-infected adults (10-30%) develops bnAbs and they only develop them after several years of infection [5]. Understanding how neutralization breadth develops during natural infection will be necessary to guide novel vaccine development strategies.

In HIV-infected individuals, bnAbs target epitopes in conserved areas of the Env including the CD4-binding site, V1V2-glycan region, and V3-glycan region on gp120; and the gp120-gp41 interface and fusion domain and membrane proximal external region (MPER) on gp41 [6]. Immunogenetic characterization of bnAbs identified in adults has demonstrated unusual traits such as high levels of somatic hypermutation (SHM), nucleotide insertions and deletions, long complementary determinant region 3 (CDR3) lengths, and restricted variable gene use [7]. Recently, studies have evaluated the presence of bnAbs in HIV-infected children. Notably, Goo et al. (2014) reported that children develop cross-clade neutralization by two years of age [8]. Moreover, Meunchhoff et al. (2016) observed that 75% of HIV-infected nonprogressor children aged ≥ five years but only 19% of chronically infected adults demonstrated neutralization breadth [9]. Altogether, these previous studies suggest that HIV-infected children develop neutralization breadth earlier and more frequently than infected adults.

Epitope mapping of neutralizing Abs in pediatric cohorts has suggested that bnAbs from children target the same key epitopes on HIV-1 Env glycoproteins as bnAbs from adults. However, in contrast to adults, where broadly neutralizing activity is typically attributed to a single or a few distinct epitope specificities [10] plasma neutralization in children appears to be polyclonal [11]. Furthermore, distribution of these bnAb specificities differed from corresponding maternal plasma, with a higher prevalence of MPER-specific Abs and decreased titers of V2-glycan specific Abs. Thus, it is possible that the mechanism by which children achieve broad plasma neutralization is distinct.

Despite these recent findings, our knowledge of the ontogeny of Env-specific Ab responses in pediatric settings remains incomplete. While several studies have investigated the ontogeny and IgG subclass distribution of HIV-specific Abs in HIV-infected adults [12], few studies have been conducted in young children [13]. Further characterization of Env-specific binding responses and IgG subclass distribution in infants and young children might guide the design of vaccines targeting the early life period.

Importantly, previous studies investigating neutralization breadth development in children have several inherent limitations such as small cohort size [8] or focus on the specific population of nonprogressor children [9]. To define the frequency and kinetics of bnAb in HIV-infected children, we acquired plasma samples from 212 ART-naïve, clade B HIV-infected children infected either *in utero* or at the time of delivery from the International Maternal, Pediatric, Adolescent AIDS Clinical Trials (IMPAACT) repository. In addition, we obtained plasma samples from 44 ART-naïve HIV chronically-infected adults from the Neutralization Serotype Discovery Project (NSDP) study [14] for comparison. Using these two cohorts, we compared the magnitude, specificity, and IgG subclass distribution of HIV-1 Env-specific Abs between HIV-infected children and adults. In addition, we compared neutralizing antibody responses in children to those of adults using a global panel of HIV-1 strains [15] and key epitopes targeted by pediatric neutralizing antibodies were identified. To our knowledge, this study represents the most comprehensive analysis of HIV neutralizing antibody responses in a large cohort of young children conducted to date.

## RESULTS

### HIV-1 Env-specific Ab binding responses in the pediatric cohort in comparison to adults

A panel of 17 HIV-1 antigens was used to assess the breath and epitope-specificity of HIV-1 Env-specific IgG in cross sectional samples of children with HIV, aged one to three years, (n=212) and in clade B chronically infected adults (n=44). Because these children acquired infection *in utero* or around birth, their age reflects the duration of infection. The breadth of the HIV-specific binding antibodies was comparable between children and adults and we observed no difference in the magnitude of binding to cross-clade specific HIV-1 gp120 and gp140 antigens between the two groups (**Figure 1**). More than 90% of children and adults had antibodies that bind to all the Env glycoproteins tested, except for A244 gp120 (Supplemental Table 1). The frequency of responders against A244 gp120 was higher in adults than in children (86 vs 70%, p=0.032), but comparable frequencies were observed for the other antigens.Interestingly, the magnitude of the IgG binding against most antigens increased between one and two years of age, but was comparable between two and three year-old children (Supplemental Figure 1). Thus, Env-specific antibody levels progressively increase during the first two years of life in perinatally infected children and, by two years of age, their antibody levels are comparable to that of chronically infected adults.

**Table 1.** Cohort Clinical Summary. Pediatric cohort samples were provided by completed IMPAACT studies ACTG 152, 300, 382, and 390. Adult clade B-infected plasma samples were provided by the Neutralization Serotype Discovery Project.

|  | Adult Cohort | Pediatric Cohort |  |  |
| --- | --- | --- | --- | --- |
| Cohort Size | n = 117 | 1-year-old | 2-year-old | 3-year-old |
|  |  | n = 91 | n = 62 | n = 59 |
| Gender | N/A | ? | ? | ? |
| Clade Infected | Multiple Clade-Infected<br>(with subset of 44 Clade B-Infected) | Clade B-Infected |  |  |
| Duration of Infection | Chronically Infected for $\geq 3$ years | Infected *in utero* or at birth | | |
| ART Exposure | ART-naïve | ART-naïve |  |  |

**Figure 1.**
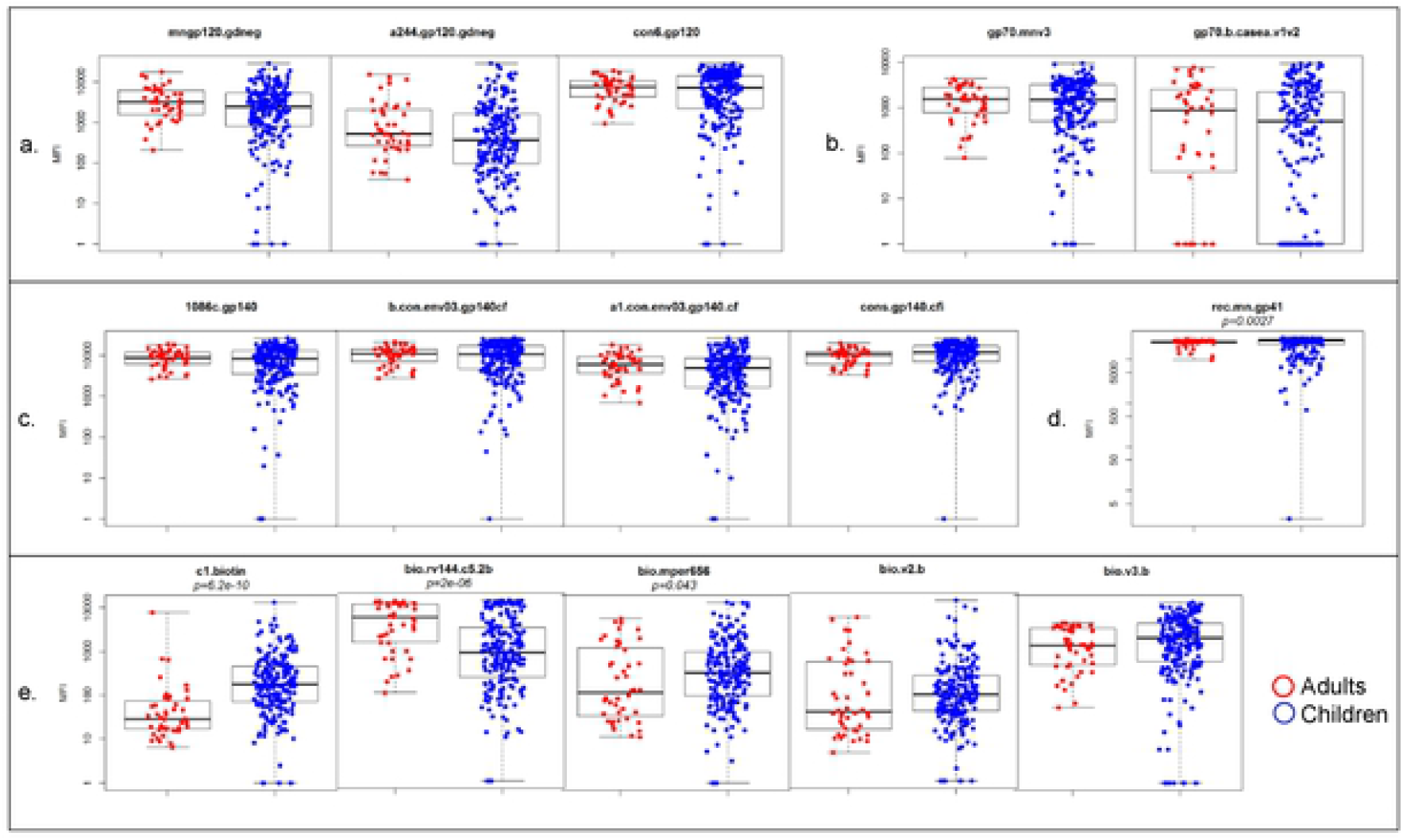
HIV-1 Env-Specific Total IgG. Total IgG for select HIV-1 Env epitopes were measured by binding antibody multiplex assay in adult and pediatric cohorts. Epitopes include gp120 **(a)**, variable loops **(b)**, gp140 (c), gp41 **(d)** and peptides **(e)**. Significant difference between adult and pediatric cohorts by Wilcoxon as noted.

Env-specific antibodies from adults and children generally bound to the same epitopes, but slight differences were observed between the two groups for some of the tested epitopes. Notably, children had higher levels of antibodies against the constant region 1 (C1) and the MPER region whereas adults had higher levels of antibodies against the constant region 5 (C5). Similarly, a higher percentage of children than adults had detectable antibodies against the MPER (79% vs 61%, p=0.017) and the C1 peptide (58% vs 11%, p<0.001) whereas the frequency of antibodies against the C5 peptide was higher in adults than in children (77% vs 42%, p<0.001). Thus, overall, HIV-infected children demonstrated robust Env-specific antibody responses that largely target the same Env regions as adult antibodies.

### HIV-1 Env-specific IgG subclass distribution in HIV-1 infected adults and children

We then measured the magnitude and frequency of Env-specific IgG1, 2, 3 and 4 in the pediatric and adult samples. Children tended to have a lower magnitude gp120-specific IgG1 responses as compared to adults (p=0.042) and both adults and children had very high magnitude of gp41-specific antibodies (**Figure 2**). Children also had lower magnitude gp41-specific IgG3 (p=0.006), and lower levels of Env glycoprotein-specific IgG4 than adults (p<0.001 for gp140, gp120 and gp41). In contrast, children had higher magnitude gp41-specific IgG2 antibodies (p=0.004) as compared to adults. Similar to total IgG, the levels of gp120-specific IgG1 increased with age, whereas low levels of antibodies from the other IgG subclass were observed across the age groups (Supplemental Figure 2).

**Figure 2.**
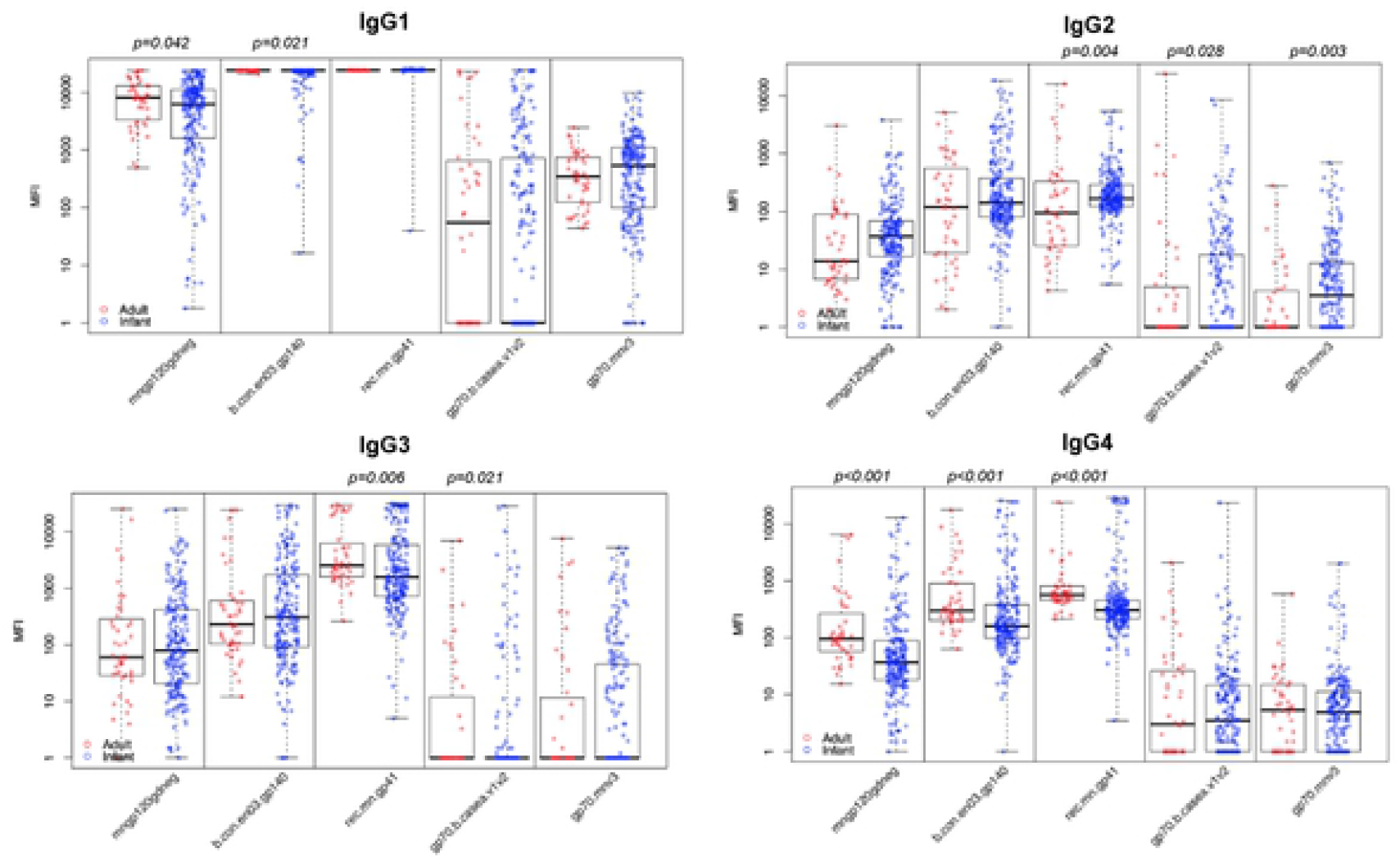
HIV-1 Env-Specific lgG Subclass. Individual lgG subclasses for select HIV1 Env eprtopes were measured by binding antibody multiplex assay in adult and pediatric cohorts. Significant difference between adult and pediatric cohorts by Wilcoxon as noted.

While the majority of adults and children had detectable levels of Env glycoprotein-specific IgG1 (94 to 100%), slight differences in the frequency of the other subclasses were observed between adults and children (Supplemental Table 2). Most adults and children had IgG3 and IgG4 against gp41 but a higher proportion of children had gp41-specific IgG2 when compared to adults (87% vs 48%, p<0.001). In contrast, IgG4 antibodies against gp120 were detected more frequently in adults than in children (48% vs 22%, p=0.002). Approximately half of the adults and children had gp120-specific IgG3 and approximately 20% had gp120-specific IgG2. The proportion of adults and infants with detectable levels of IgG subclass antibodies against the variable loop 2 (V2) and the variable loop 3 (V3) was comparable.

**Table 2.** Frequency of neutralization response in adults vs. children. Proportion of adult and pediatric samples with 1D50>100 for eaeh virus. The pediatric cohort demonstrated comparable or superior neutralization frequency to adults in 9/10 viruses tested. The adult cohort had higher neutralization frequency against only one virus, X1632. Statisttcally significant pvatues included as detennined by Fisher test

| Virus | Clade | Adults (%) | Children (%) |  |
| --- | --- | --- | --- | --- |
| 25710 | C | 91 | 90 |  |
| TRO11 | B | 68 | 78 | p=0.048 |
| X2278 | B | 75 | 86 | p=0.015 |
| BJOX2000 | CRF07_BC | 60 | 55 |  |
| X1632 | G | 63 | 28 | p<0.001 |
| CE1176 | C | 62 | 51 |  |
| 246F3 | AC | 48 | 71 | p<0.001 |
| CH119 | CRF07_BC | 67 | 88 | p<0.001 |
| CE0217 | C | 46 | 40 |  |
| CNE55 | CRF01_AE | 34 | 41 |  |

### HIV neutralization responses in HIV-infected children as compared to infected adults

The ability of pediatric samples to mediate broad neutralization was assessed against a panel of 10 HIV-1 pseudoviruses from the global neutralization panel. This panel was selected from 219 pseudoviruses to be representative of the global diversity of circulating HIV-1 strains [15]. The neutralization potency and breadth in children was compared with previously reported data of 117 chronically infected adults from the NSDP [14]. Overall 69% of the children and 68% of adults neutralized at least 50% of the viruses in the panel with an ID50 ≥50. The frequency of broad neutralization slightly increased with age, as only 60% of one-year old children but 76% of two-year old children were able to neutralize 50% of the viruses. Comparable percentage of two-year old and three-year old children (76% vs 75%) neutralized 50% of the viruses with an ID50 ≥50. Using a more stringent ID50 cutoff of 100, we observed that 26% of children but only 19% of adults neutralized 50% of the viruses with an ID50 ≥100. The percentage of children neutralizing 50% with ID50 >100 was comparable across the age groups (23% for one-year old, 29% for two and three-year old). There was no statistical difference in the percentage of children and adults that were able to neutralize 5/10 viruses (**Table 2**). A higher percentage of children than adults were able to neutralize 4/10 viruses (HIV TRO11: 78% vs 68%, p=0.048; HIV X2278: 86% vs 75%, p=0.015; HIV 246F3: 71% vs 48%, p<0.001; HIV CH119: 88% vs 67%, p<0.001), whereas a higher percentage of adults neutralized one virus (HIV X1632: 63% vs 28%, p<0.001). Thus, overall, the pediatric cohort demonstrated comparable or superior neutralization frequency as compared to adults for 9/10 viruses tested. Children also demonstrated comparable or superior neutralization potency (**Figure 3)**. The median neutralization titer was higher in children than in adults for 4/10 viruses tested whereas adult samples demonstrated higher neutralization potency against one virus.

**Figure 3.**
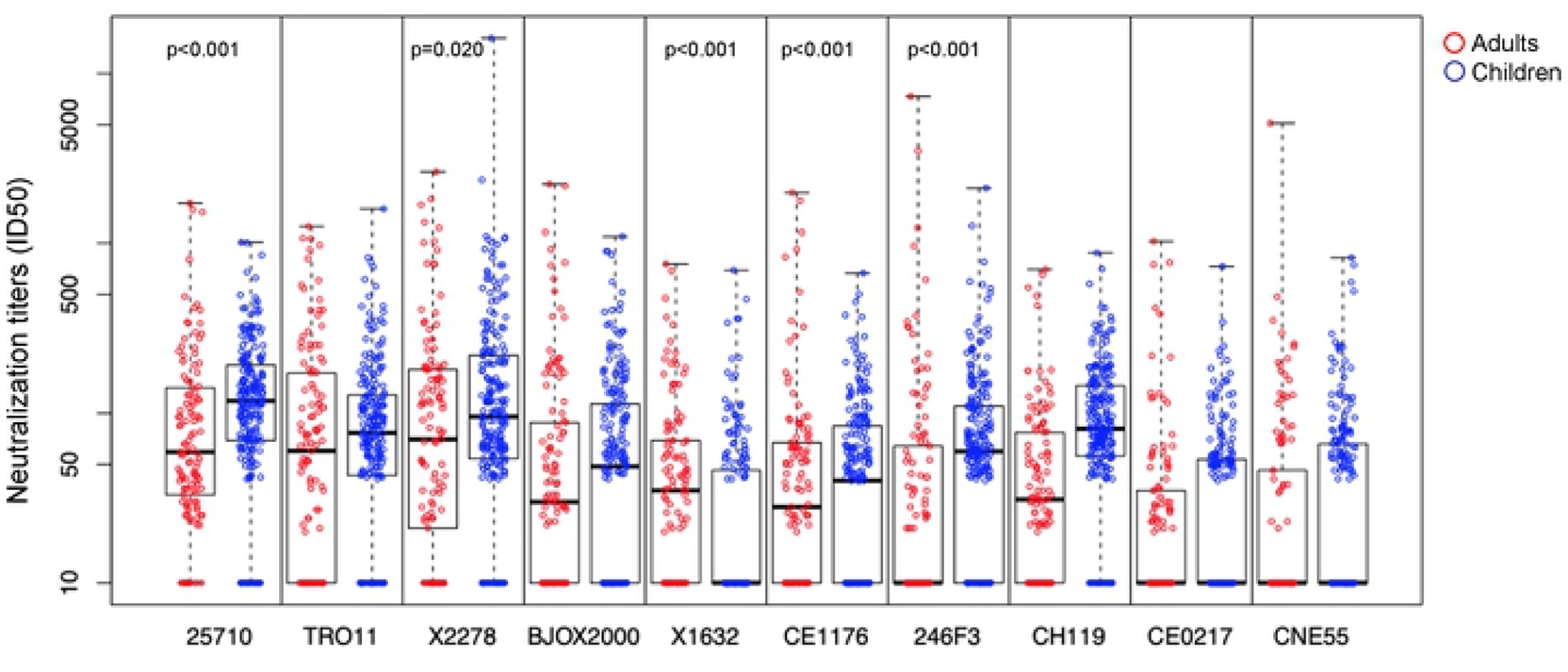
Neutralization potency in adults vs. children. The neutralization liters of the pediatric samples were calculated for each virus and compared to data from the historical adult cohort. Children demonstrated higher neutralization potency against 4/10 viruses and adults demonstrated higher neutralization potency against 1/10 viruses. Statistical significance determined by Wilcoxon test as noted.

To assess the totality of the virus neutralization activity, neutralization scores were generated for each pediatric and adult sample. The neutralization score was defined as the average area under the magnitude-breadth curve (AUC) of the neutralization curve for all tested viruses. Overall, the neutralization score was higher in children than in adults (p=0.014, **Figure 4a**). As with other antibody measurements, the neutralization score increased with age across the one, two, and three-year old age groups (p=0.014, **Figure 4b**), but importantly, the neutralization score in one-year old children was comparable to that of adults (p=0.44, **Figure 4c**). Thus, by one year of age, the plasma neutralization activity in perinatally infected children is comparable to that of chronically infected adults.

**Figure 4.**
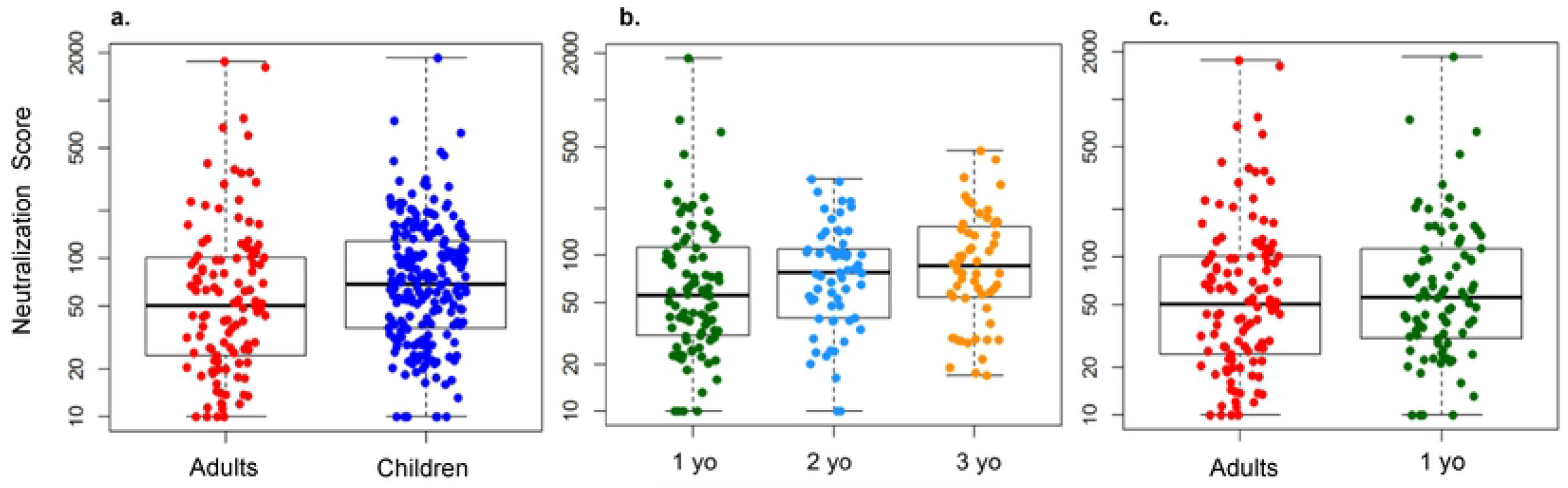
Neutralization score in adults vs. children. Neutralization score was determined as the AUC of the neutralization curve for all tested viruses (**a**) Overall, the neutralization score was greater in children than in chronically infected adults (p=0.014. Wilcoxon). (**b**) Neutralization score increased with age (p=0.014, median regression analysis) (**c**) The neutralization score of the 1 year-old cohort was comparable to that of adults (p = 0.44, Wilcoxon).

Because the neutralization score is influenced both by the proportion of the viruses neutralized as well as the potency with which these viruses are neutralized, magnitude-breadth curves were generated to determine the relative contribution of breadth and potency on the superior neutralization breadth score of children (**Figure 5**). The average ID50 of the pediatric cohort was significantly greater than that of the adult cohort with a similar proportion of viruses neutralized, indicating that the higher neutralization score observed in children is mostly driven by a superior neutralization potency with comparable number of viruses neutralized.

**Figure 5.**
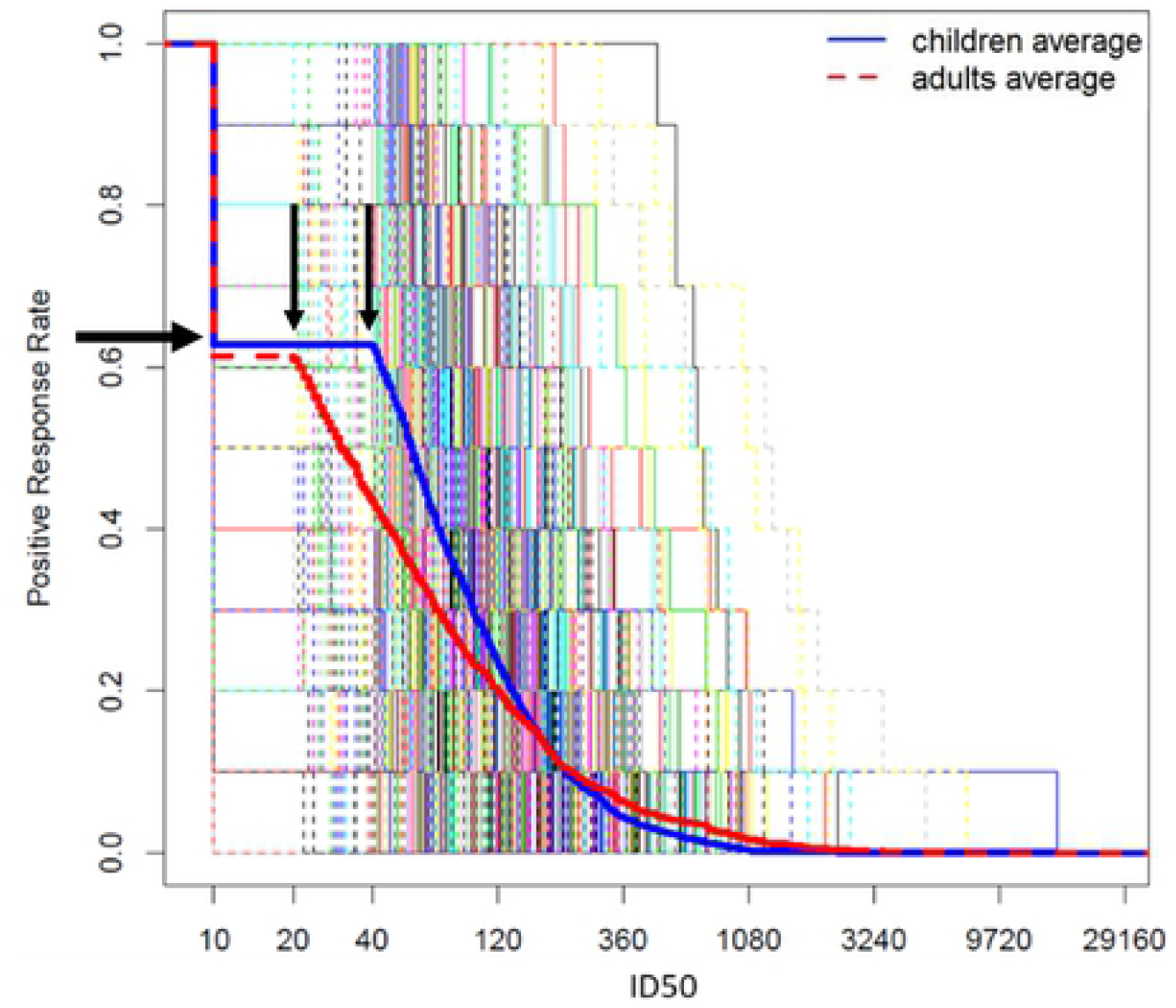
Magnitude-breadth curve comparing adults vs. children. The average ID50 value for the pediatric cohort was significantly greater than that of the adult cohort (p =0.013, Mann Whitney U) as indicated by the vertical arrows whereas a similar proportion of viruses were neutralized in the two cohorts as indicated by the horizontal arrow.

### Association between neutralization breadth and clinical factors in HIV-infected children

To explore the potential association between neutralization breadth and patient clinical factors, we performed linear regression analyses between pediatric neutralization breadth scores and patient CD4 T cell percentage and absolute counts provided by the IMPAACT repository (Supplemental Figure 4). We found a weakly positive association between neutralization breadth score and CD4+ T cell percentage (slope=0.01, p=0.002) as well as with CD4+ T cell counts (slope=0.07, p=0.003). We also explored the association between viral load and neutralization breadth and we did not observe a significant association (slope 0.10, p=0.28). However, this analysis was limited by the small number of children with available viral load in the IMPAACT database (n=15). The impact of other clinical factors such the timing of infection (intrauterine versus perinatal) could not be evaluated due to the paucity of information in the IMPAACT database. Nevertheless, our results suggest that factors other than CD4 T cell could likely contribute to drive neutralization breadth development in children.

We also examined associations between neutralization and binding antibody responses (supplemental table 3). We observed a weak, but statistically significant correlation between total gp120 and gp140 IgG and neutralization score. Moreover, gp140 IgG4 and gp41 IgG4 levels were weakly associated with neutralization score. There was no association between antibody specificity that differ between adults and children (such as C1 or C5-specific IgG) and neutralization score. Overall this data suggest a possible weak association between IgG subclass distribution and breadth development in young children.

**Table 3.** Epitope specificity of broadly neutralizing pediatric samples. Neutralization of mutant variants in the 5 pediatric samples that neutralized ≤5 of the 10-virus global panel with an ID50 ≥100 and demonstrated at least a 3-fold reduction in ID50 against one mutant pseudovirus.

|  |  |  | Epitope targeted by mutant |  |  |  |  |  |  |  |  |  | putative epitope specificity |
| --- | --- | --- | --- | --- | --- | --- | --- | --- | --- | --- | --- | --- | --- |
|  |  |  | CD4bs | CD4bs | CD4bs | CD4bs | 2G12 Sensitive | V3 Glycan | V2 Glycan | gp120-gp41 (35022) | gp120-gp41 (35022) | gp120-gp41 (PGT151) |  |
| PTD | Age | WT ID50 | N279A | N280D | G458Y | N276Q | N295V | N332A | N160K | N88A | N625A | N611A |  |
| 1 | 1 | 331 | 1550 | 2531 | 2428 | <50 | NA | 476 | 474 | 628 | 745 | NA | CD4 bs |
| 2 | 1 | 1268 | 2111 | 3516 | 2108 | 754 | 1100 | 1083 | 370 | 1444 | 928 | 923 | V2 glycan |
| 3 | 2 | 415 | NA | NA | NA | NA | 171 | 80 | 362 | 322 | 338 | 328 | V3 glycan |
| 4 | 3 | 471 | 1486 | 5411 | 938 | 196 | 167 | 120 | 605 | 397 | 260 | 427 | V3 glycan |
| 5 | 3 | 890 | 1571 | 4097 | 1223 | 503 | 1954 | 115 | 1098 | 1420 | 677 | 972 | V3 glycan |

### Epitope specificity of neutralizing antibodies in HIV infected children

Pediatric samples that were able to neutralize ≥ 5 viruses with an ID50≥100 (n=38) were utilized to map the epitope specificity of neutralizing antibodies. Neutralization assays were performed against a virus from the global panel that the plasma sample neutralized potently (either HIV TRO11, 25710 or BJOX002) and against a panel HIV-1 pseudoviruses with selective mutations to abrogate the neutralization potency of a specific class of broadly neutralizing antibodies [16] (Supplemental Table 3). In addition, the samples were tested against a mutant pseudovirus TRO11.W672A to assess the presence of MPER-specific antibodies. A total of 5/38 plasma samples demonstrated at least a 3-fold reduction in ID50 against one mutant as compared to the parent virus (**Table 4**). Three samples demonstrated substantial reduction of neutralization against a mutant that abrogates activity of V3 glycan-specific bnAbs, whereas one sample demonstrated reduced neutralization of a mutant that abrogates activity of CD4 binding site bnAbs, and the last showed reduced activity against a mutant that abrogates the activity of V2 glycan-specific bnAbs. None of the plasma samples demonstrated a dominant MPER reactivity. Thus, the epitope specificity of the neutralizing antibodies could not be mapped in 33/38 HIV-infected children with strong bnAb activity.

### Association between binding antibody responses and neutralization epitope specificity

We conducted analysis to determine if there is an association between the levels of epitope-specific binding antibodies and the neutralization specificity. PTD 2 (from Table 4) in which neutralization specificity mapped to V2-glycan epitopes demonstrated moderately high V1V2-specific IgG and IgG1 response ranking at the 65% and 56% percentile in the distribution of these responses. No V1V2-specific IgG2, IgG3 or IgG4 was detected in this child (supplemental Figure 5). The three children (PTD 3-5) in which neutralization specificity mapped to V3 glycan dependent epitopes demonstrated high V3-specific IgG binding ranking at the 75, 85, and 83% percentile of the IgG distribution. They also demonstrated high V3-specific IgG1 ranking at the 61, 79, and 75% percentile for the IgG1 distribution, whereas the levels of the other IgG subclass were variable. Thus, while high levels of V1V2 or V3-specific IgG or IgG 1 antibodies are not indicative of V2 or V3-specific neutralizing antibody responses, when children exhibited neutralizing antibodies to a particular site they tended to have high binding antibodies to the same site.

## DISCUSSION

While induction of a broad neutralizing antibody response is a critical goal for HIV vaccine, current immunization approaches have failed to achieve this goal [17]. Recent reports indicating that: 1) HIV vaccination can elicit more robust and durable antibody responses in infants than in adults [18] [19]; and 2) that children may develop broad neutralization more frequently and earlier than adults [9], suggest that the early life immune system may present advantages to overcome the unique immunologic challenges posed by the induction of neutralization breadth through vaccination. The purpose of this study was to assess the presence and characteristics of broadly neutralizing antibodies in a large cohort of HIV-infected, ART-naïve young children and compare the kinetics of the development of bnAb responses after infection to that of chronically-infected adults. Our results are in concordance with prior reports indicating that children are able to develop broad neutralizing antibody responses and, importantly, that the mechanism of breadth development may differ between adults and children [8, 9].

Only a few studies have assessed broad neutralizing antibody responses in HIV infected children. Notably Goo et al, investigated broad neutralization in a small cohort (n=28) of clade A infected children from Kenya and reported that broad neutralization could be detected in infants as early as one year after infection [8]. Subsequently, Muenchhoff et al., investigated broad neutralization in a cohort of children from South Africa (aged >5 years) and reported higher frequency of broad neutralization in children as compared to chronically infected adults [9]. In contrast to these previous studies, our study focused on a large cohort of clade B HIV-infected children. Moreover, we focused on children aged 1 to 3 to specifically define the kinetics of the development of broad neutralization in children, as HIV-infected adults usually develop neutralization breath after several years of infection [44]. This age range excluded the possibility that passive maternal antibodies contributed to the measured neutralization as before one year of age it would be difficult to decipher the contribution of maternal versus infant antibodies. The uniqueness of this historical set of pediatric ART naïve samples is worth noting as the WHO now recommends initiation of ART at the time of diagnosis. To assess neutralization breadth, we used viruses from the global neutralization panel [18]. We observed that one year old HIV-infected children can demonstrate broad neutralization, corroborating the findings of Goo et al [8] that HIV-neutralization can develop rapidly in early life. Yet, in contrast to the findings from Muenchhoff et al., (2016) who reported more frequent bnAb development in children than in adults (75% vs 19%, p<0.0001) [9], we found that the proportion of children and adults who neutralized 50% of the viruses was comparable (69% vs 68%). Comparable frequencies between the groups were still observed when we only considered children and adults who neutralize 50% of viruses with ID50 >100 (26% vs 19%, p=0.13). Interestingly, Makhdoomi et al., reported that the neutralization breadth in HIV-infected children from India aged 5 to 17 years [45] increased over time. In our study, although we did not test longitudinal samples, we compared the neutralization activity across the children’s age groups. We found that the neutralization scores increased with age (Fig 4), but by one year of age, the breadth potency neutralization score was comparable to that of chronically infected adults. Thus, the fact that our cohort was younger than that of Muenchhoff et al., may contribute to explain the differences in observed results. Furthermore, whereas our cohort was US-based (predominant clade B infections), their cohort was South African based (predominant clade C infections). The South African cohort also focused on the specific subpopulation of nonprogressors [9] and the virus panels used in the two studies to assess neutralization breadth were different. Finally, it is possible that differences in transmission modes between the two cohorts contributed to the slightly divergent results [20]. In our US-based cohort, transmission occurred in utero and perinatally, whereas breast milk transmission is also an important mode of transmission in South Africa. Despite these differences, it is worth noting that our results combined with those of previous studies establish that children are able to rapidly develop broad neutralizing antibody responses after HIV infection.

Several studies have attempted to understand immune differences between children and adults, with particular focus on the first 12 months of life and the development of B cell populations [21-26]. However, although several studies have investigated the kinetics of HIV-specific binding antibodies in adults, less is known about pediatric HIV-specific antibody responses. We found that although the magnitude of HIV-specific binding Ab responses against key Env epitopes was lower in one-year-old children, by two years of age it was comparable to that of adults (Supplemental Figure 1). While we found no difference in binding breadth between adults and children, we observed some slight differences in IgG binding epitope specificities between the two groups. Notably, MPER and C1-specific antibodies were higher in children than in adults whereas C5-specific antibodies were higher in adults. It has also been previously reported that infants develop antibodies against the HIV gp160 precursor glycoprotein first, in contrast to adults who first develop anti-gp41 antibodies, which suggests possible differences in the kinetics of HIV-specific Abs between these two populations [13, 27]. Overall, the difference in epitope specificities between adults and children suggests that similar immunogens could elicit distinct responses in these two populations, an observation that may be relevant for vaccine development and monitoring.

Previous studies indicated that increased levels of HIV-specific IgG2 and IgG4 during early HIV infection in adults correlate with development of broad neutralization [28]. Moreover, an association between high levels of IgG3 and neutralization have been reported [29]. We therefore investigated if a distinct IgG subclass distribution in children as compared to adults may contribute to explain the early development of broad neutralization. Overall, we found that the IgG subclass profiles of the pediatric and adult cohorts were grossly similar with only a few statistically significant differences. Most notably, we observed that the majority of children mounted a detectable gp41-specific IgG2 response, whereas this response was seen in less than half of chronically infected adults. The clinical significance of this difference is unclear as there was no association between this response and neutralization breadth. The ability of children to produce IgG2 is usually delayed compared to other subclasses [30], thus the observation that young children develop more robust IgG2 responses than adults is somewhat surprising. Nevertheless, as an association between low levels of gp41-specific IgG2 are usually associated to later stage disease [31], the higher levels of gp41-specific IgG2 in children may simply be a marker of a more recent infection. Interestingly, it was recently reported that IgG3 enhances the neutralization activity of a HIV V2-specific bnAb [16]. Thus, we were interested to define if higher HIV-specific IgG3 were observed in children. For most antigens, there was no difference in the magnitude of IgG3 responses between adults and children, except for gp41 and V1V2 that were higher in adults than in children. Thus, overall, our results suggest that IgG subclass distribution may not contribute to the early development of broad neutralization in children.

Previous studies have suggested that in contrast to adults in which a single epitope specificity frequently mediate neutralization breadth [32, 33], plasma broad neutralization in children may be mediated by a polyclonal response. For example, Goo et al, were not able to map the epitope specificity of broad neutralization in the majority of the children from their cohort suggesting that either the pediatric neutralizing antibodies target novel neutralization epitopes that are distinct from adults or broad neutralization is mediated by polyclonal antibodies. Ditse et al, [11] analyzed the epitope specificity of neutralizing antibodies in 16 clade C infected nonprogressor children from South Africa. They observed that the majority of children had antibodies targeting multiple known neutralization epitopes, and that most children had antibodies against V2 and V3-glycan dependent epitopes whereas MPER antibodies rarely confer breadth to children. Similarly, Mishra et al [46] reported that plasma bnAbs targeting the V2-apex were predominant in a small cohort of 10 elite neutralizers infants from India. In their study, only two children demonstrated responses against multiple neutralizing epitopes. In contrast to these previous studies, we were only able to map neutralization to a dominant epitope in five out of 38 samples tested. In three of these, broad neutralization was mediated by V3-glycan dependent antibodies, whereas V2-glycan dependent and CD4 binding site antibody were the dominant neutralizing epitopes in the other two samples, respectively. While it is possible that our selection of mutant pseudoviruses did not capture an existing dominant epitope, the most common bnAb targets-V2 glycan, V3 glycan, CD4bs, MPER and gp120-gp41 interface [11]-were represented in our panel, and thus it is likely that the fact that we could not map the dominant specificity in 33/38 samples indicate that breadth was mediated either by a polyclonal response or that these pediatric nAbs target a novel neutralization epitope. A dominant polyclonal response in children may indicate that the mechanisms by which the pediatric and adult immune systems generate plasma broad neutralization could differ. In adults, it has been proposed that the development of bnAbs results from cooperation between multiple B cell lineages; “cooperative” lineages react to and select for escape mutants, which allows the bnAb lineage to be continually stimulated by the founder virus to the extent of sustained affinity maturation [7]. Our findings may suggest that the abundance of naïve B cells in children allows for activation of multiple independent B cell lineages upon HIV infection leading to a polyclonal response. Thus, plasma broad neutralization in children could be mediated by the additive activity of a collection of antibodies with limited to moderate neutralization.

Elucidating the mechanisms through which children achieve broad neutralization will be critical to guide vaccine development. Importantly, analysis of two V3-glycan dependent bnAbs isolated from children indicated that breadth was acquired through a pathway distinct from that of adult antibodies from the same class [34, 35] with notably lower levels of somatic hypermutation and early accumulation of critical improbable mutations [36]. Identifying and characterizing more Abs from children is imperative to determine if low levels of SHM is a consistent unique feature of pediatric nAbs. This knowledge will ultimately guide HIV vaccine strategies aiming at inducing broad neutralization. Such strategies could involve initiation of vaccination early in life (following the immunization schedule for the under 5 year olds) and boosting through pre-adolescence in order to achieve protective immunity prior to sexual debut.

## MATERIALS AND METHODS

### Samples

Pediatric plasma samples from ART-naïve children aged one, two and three-years old (n=212) who were enrolled in the completed IMPAACT studies ACTG 152, 300, 382 and 390 [37-40] were obtained from the IMPAACT biospecimen repository. All these children were assumed to be infected with clade B HIV-1 at birth or *in utero* as they were born tp women who acquired HIV in the US. Adult plasma samples (n=44) were obtained from ART-naïve adults with chronic clade B HIV-1 infection of ≥ three years who participated in the Neutralization Serotype Discovery Project [14]. A summary of the clinical characteristics of the included patients in provided in Table 1.

### Binding antibody multiplex assay for the *measurement of Env-specific IgG*

A previously described binding antibody multiplex assay (BAMA) [27] was used to measure IgG binding to a panel of 17 HIV-1 antigens. These included: 1) cross-clade gp120s (Con6 gp120/B, MNgp120, and A244 gp120); 2) cross-clade gp140s (B.con_env03 gp140, ConS gp140 CFI, A1.Con gp140, and 1086C gp140); 3) gp70 scaffold Env constructs (gp70_B.CaseA_V1V2, gp70 MNV3); 4) a clade B recombinant gp41 (RecMN gp41); 5) peptides representing the V2 (Bio-V2.B), the V3 (Bio-V3.B), the MPER (MPER656) the C5 (RV144 C5.2B) and the C1 (C1 Biotin) regions of the HIV Env; and finally 6) YU2 Core, YU2 Core D368R mutant to assess antibodies against the CD4 binding site. A pilot study was conducted to determine the optimal testing dilution for the samples. This dilution was 1:100 dilutions for all antigens except for the gp140s, Bio-V3B and recMNgp41 that were tested at 1:2000. Total antigen-specific IgG was then detected with a mouse anti-human IgG phycoerythrin-conjugated Ab (Southern Biotech) at 2 μg/mL. Binding was measured using a Bio-Plex 200 instrument (Bio-Rad Laboratories, Inc.). HIV-1 human hyperimmune immunoglobulin (HIVIG) derived from the plasma of HIV-1-infected donors was used as a positive control and normal human serum was used as a negative control. IgG responses were expressed as mean fluorescence intensity (MFI). All MFI values were adjusted for nonspecific binding by subtracting the MFI of blank beads, except for gp70_B.CaseA_V1V2 and gp70 MNV3, which were adjusted by subtracting the MFI of beads coupled with a control gp70 construct (MuLVgp70). An HIV Env-specific Ab response was considered positive if it had MFI values above a positivity cutoff determined as highest of either the mean plus 3 standard deviations of the MFI of a panel of 20-30 HIV negative plasma, or the lower detection limit of 100 MFI. To ensure consistency between assays, 50% effective concentration and maximum MFI values of HIVIG control were tracked by Levey-Jennings charts [41].

### Binding antibody multiplex assay for the *measurement of IgG subclasses*

Measurement of IgG subclass-specific HIV-1 binding Ab response was conducted as described above with some modifications. All plasma samples were tested against a limited panel of 5 HIV-1 antigens: B.con_env03 gp140, MNgp120, gp70_B.CaseA_V1V2, gp70 MNV3, and Rec MN gp41 at a 1:50 dilution. Antigen-specific subclass response was detected using biotin-conjugated mouse anti-human IgG1 (BD Pharmingen, 4μg/L), IgG2 (Southern Biotech, 5ug/mL), IgG3 (Calbiochem, 2μg/mL), or IgG4 (BD Pharmingen, 2μg/mL) and tertiary detection agent streptavidin-phycoerythrin (BD Biosciences) at 5 μg/mL.

### Neutralization Assays

Neutralization was measured as the ability of plasma samples to reduce virus infection of TZM-bl cells as previously described [42] against a panel of 10 viruses from the global neutralization panel (deCamp et al., 2014) [15] (Table 2). Briefly, plasma samples were incubated with an HIV-1 pseudovirus for 45-90 min at 37°C, then TZM-bl cells were added and plates were incubated for 48 hr. A luciferase substrate (Bright-Glo; Promega) was added, and luminescence was measured. Results were reported as the 50% inhibitory dilution (ID50), which is the dilution of plasma resulting in 50% reduction in luminescence compared to that of virus control wells.

### Mapping of Neutralizing Epitopes

Plasma neutralization was assayed against a mutant panel of HIV-1 strains BJOX002, 25710, or TRO.11 as described above. Panel choice was dependent on which parent virus was most potently neutralized by the plasma sample. Each panel consisted of 6-11 HIV-1 Env epitope knockout pseudoviruses (Supplemental Table 4). Significant reductions in ID50 between parent and knockout virus suggests that epitope contributes to the neutralization ability of that sample. In addition, samples were tested for MPER reactivity using TRO.11.W672A pseudotyped virus.

### Statistical analysis

All statistical analyses were performed in the R statistical computing and graphics environment. For two sample tests we used Mann-Whitney rank-based tests [47]. Neutralization scores were computed as the area under the magnitude-breadth (M-B) curve, which describes the magnitude (NAb titer) and breadth (number of isolates neutralized) of an individual plasma sample assayed against all tested viruses, and equals the average log10 titer over the targets [48]. All p-values are two-sided. Boxplots were used to graphically display distributions of log10 NAb titers to individual isolates. Positive response rates were compared between groups by Fisher’s exact tests [49]. Median regression was conducted using the quantreg package [50].

Titers of NAbs among responders were compared between groups by 95% CIs about the ratio of GMTs. Equality of the overall distribution of log10 NAb titers between two groups was tested as described (46), using 10,000 permutated data-sets to compute a p-value. The false discovery rate (FDR) was used to determine tests that remained statistically significant after adjustment for the multiple hypothesis tests. The FDR method was performed at level 0.05. Assessment of magnitude and breadth of neutralization of a panel of isolates.

## Acknowledgments

Research reported in this publication was supported by The National Institute of Allergy and Infectious Diseases (NIAID) of the National Institutes of Health under award number R01AI143370. Part of the results resulted from research supported by the Duke University Center for AIDS Research (CFAR), an NIH funded program (5P30 AI064518). The analysis was partially supported by the NIH award R01AI122991 to YF. Overall support for the International Maternal Pediatric Adolescent AIDS Clinical Trials Network (IMPAACT) was provided by the National Institute of Allergy and Infectious Diseases (NIAID) with co-funding from the Eunice Kennedy Shriver National Institute of Child Health and Human Development (NICHD) and the National Institute of Mental Health (NIMH), all components of the National Institutes of Health (NIH), under Award Numbers UM1AI068632-15 (IMPAACT LOC), UM1AI068616-15 (IMPAACT SDMC) and UM1AI106716-15 (IMPAACT LC), and by NICHD contract number HHSN275201800001I. The content is solely the responsibility of the authors and does not necessarily represent the official views of the NIH. We are grateful to the clinical teams and study participants of ACTG 152, 300, 382 and 390.

## Supplemental Figures

**Supplemental Table 1.** Frequency of Env-specific IgG binding response in adults vs. children. Proportion of adult and pediatric samples with binding magnitude of MFI >100 for each antigen, as assessed by BAMA. Statistically significant p--values included as determined by Barnard’s test.

| Antigen | Adults (%) | Children (%) |  |
| --- | --- | --- | --- |
| Con6 gp120/B | 100 | 96 |  |
| MNgp120 | 100 | 92 |  |
| A244 gp120 | 86 | 70 | p=0.032 |
| B.con_env03 gp140 CFI | 100 | 100 |  |
| ConS gp140 CFI | 100 | 100 |  |
| A1.Con gp140 | 100 | 96 |  |
| 1086C gp140 | 100 | 98 |  |
| RecMN gp41 | 100 | 100 |  |
| gp70_B.CaseA_V1V2 | 73 | 67 |  |
| gp70 MNV3 | 98 | 90 |  |
| Bio-V2.B | 39 | 53 |  |
| Bio-V3.B | 95 | 93 |  |
| MPER656 | 61 | 79 | p=0.017 |
| RV144 C5.2B | 77 | 42 | p<0.001 |
| C1 Biotin | 11 | 58 | p<0.001 |
| YU2 Core | 98 | 92 |  |
| YU2 Core D368R | 23 | 36 |  |

**Supplemental Figure 1.**
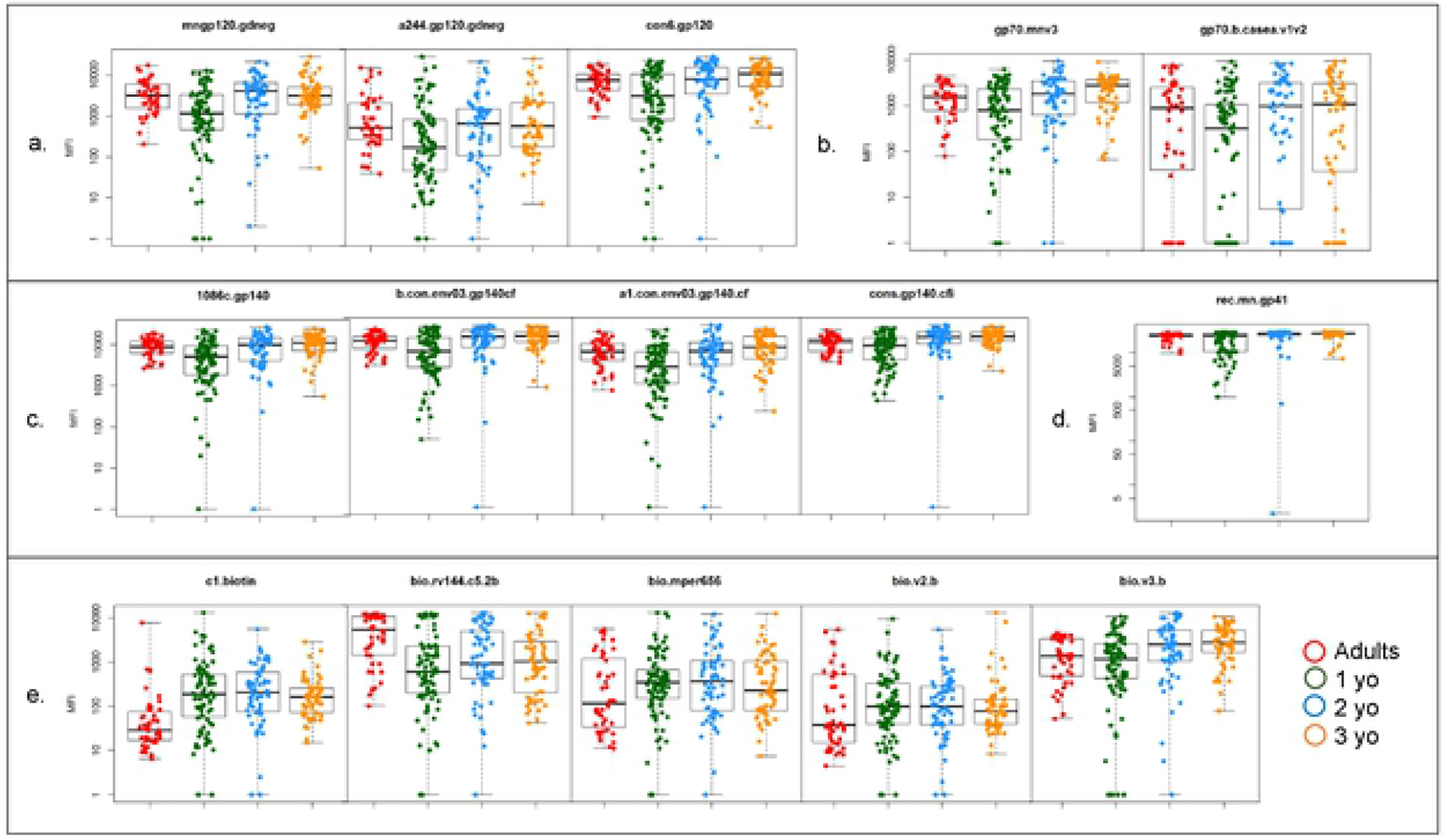
HIV-1 Env-Specific Total IgG Extended. Tolal IgG for select HIV-1 Env epitopes were measured by binding antibody multiplex, assay in adult and pediatric cohorts,including age 1, 2, and 3-year old sub-cohorts shown here. Epitopes include gp120 (**a**), variable loops (**b**), gp140 (**c**), gp41 (**d**) and peptides **(e)**.

**Supplemental Figure 2.**
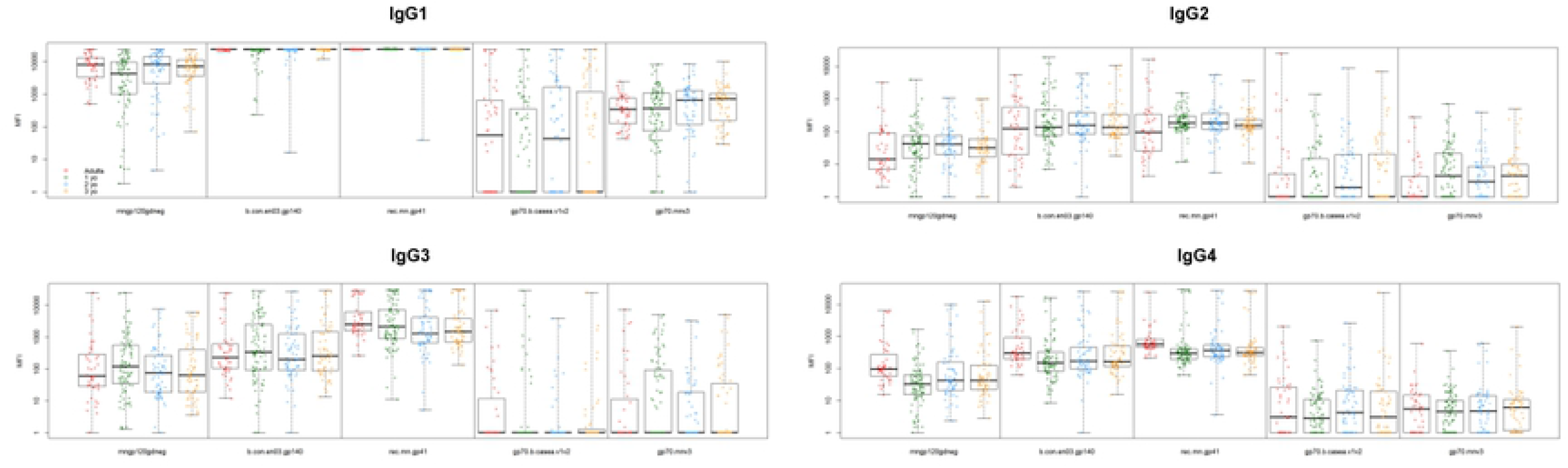
HIV1 Env-Specific IgG Subclass Extended. Individual IgG subclasses for select HIV-1 Env epitopes were measured by binding antibody multiplex assay in adult and pediatric cohorts, including age 1, 2, and 3-year old sub-cohorts shown here

**Supplemental Table 2.** Frequency of Env-specific IgG subclass binding response in adults vs. children. Proportion of adult and pediatric samples with binding magnitude of MFI > 100 for each antigen, as assessed by BAMA, Statistically significant p-values included as determined by Barnard’s test.

| IgG Subclass | Antigen | Adults (%) | Children (%) |  |
| --- | --- | --- | --- | --- |
| IgG1 | MNgp120 | 100 | 94 |  |
|  | B.con_env03 gp140 CFI | 100 | 100 |  |
|  | RecMN gp41 | 100 | 100 |  |
|  | gp70_B.CaseA_V1V2 | 45 | 38 |  |
|  | gp70 MNV3 | 80 | 75 |  |
| IgG2 | MNgp120 | 20 | 21 |  |
|  | B.con_env03 gp140 CFI | 55 | 64 |  |
|  | RecMN gp41 | 48 | 87 | p<0.001 |
|  | gp70_B.CaseA_V1V2 | 11 | 8 |  |
|  | gp70 MNV3 | 5 | 5 |  |
| IgG3 | MNgp120 | 43 | 46 |  |
|  | B.con_env03 gp140 CFI | 77 | 71 |  |
|  | RecMN gp41 | 100 | 98 |  |
|  | gp70_B.CaseA_V1V2 | 16 | 8 |  |
|  | gp70 MNV3 | 16 | 19 |  |
| IgG4 | MNgp120 | 48 | 22 | p=0.002 |
|  | B.con_env03 gp140 CFI | 98 | 73 | p=0.002 |
|  | RecMN gp41 | 100 | 96 |  |
|  | gp70_B.CaseA_V1V2 | 14 | 7 |  |
|  | gp70 MNV3 | 2 | 5 |  |

**Supplemental Figure 4.**
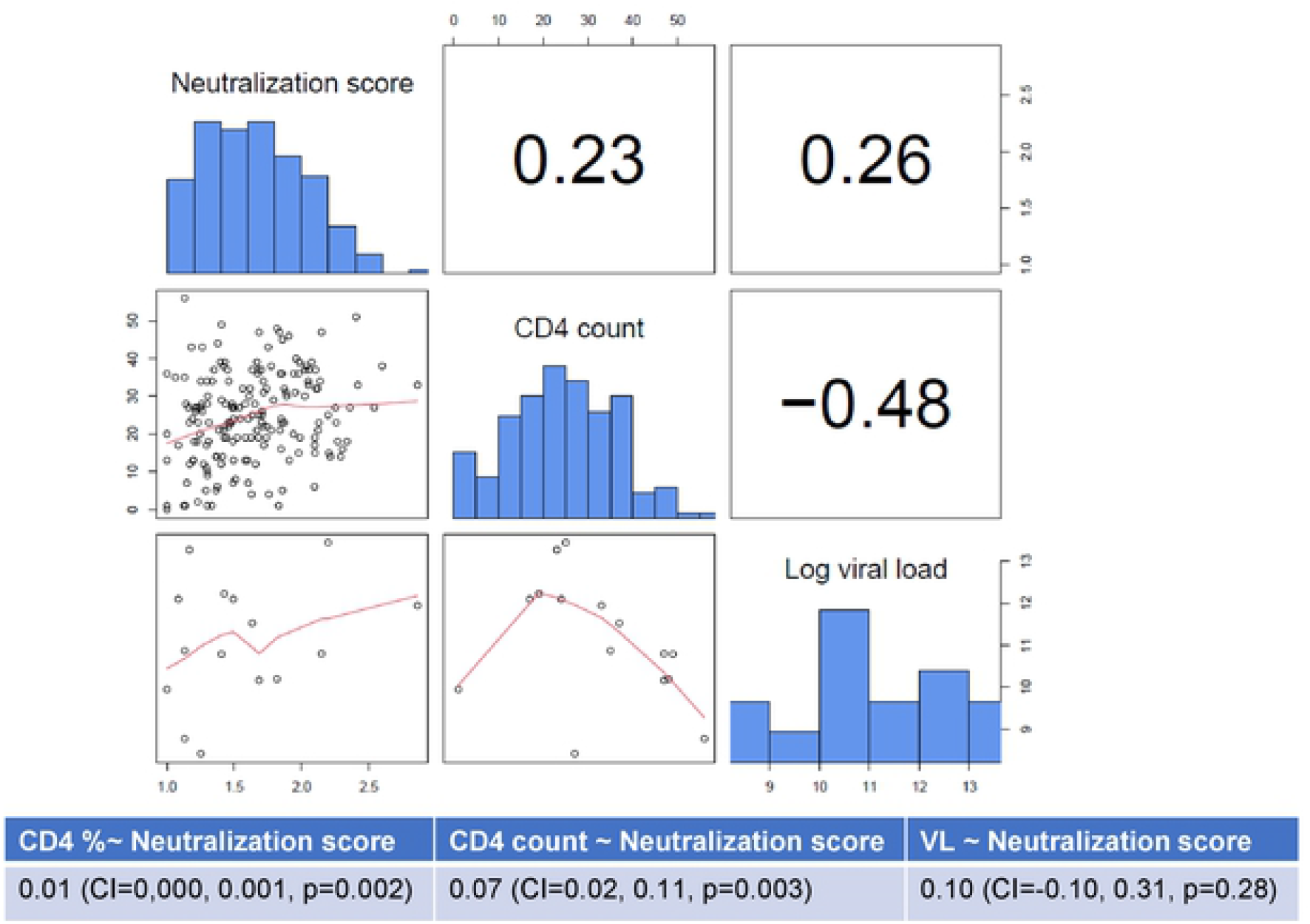
Association between neutralization breadth score and clinical factors for pediatric cohort. Estimated slopes and p-values from linear regression are shown below.

**Supplementary Table 3:** association between binding and neutralizing antibody responses.

|  | Spearman correlation with<br>neutralization score | P value | Holm adjusted p<br>value |
| --- | --- | --- | --- |
| IgG MN gp120 | 0.205 | **0.003** | **0.039** |
| IgG1 MN gp120 | 0.181 | **0.008** | 0.101 |
| IgG2 MN gp120 | 0.141 | **0.041** | 0.407 |
| IgG3 MN gp120 | -0.048 | 0.490 | 1.000 |
| IgG4 MN gp120 | 0.187 | **0.006** | 0.082 |
| IgG B con gp140 | 0.261 | **0.000** | **0.002** |
| IgG1 B con<br>gp140 | 0.147 | **0.032** | 0.354 |
| IgG2 B con<br>gp140 | 0.085 | 0.215 | 1.000 |
| IgG3 B con<br>gp140 | 0.010 | 0.889 | 1.000 |
| IgG4 B con<br>gp140 | 0.208 | **0.002** | **0.034** |
| IgG gp41 | 0.136 | **0.049** | 0.437 |
| IgG1 gp41 | 0.100 | 0.148 | 1.000 |
| IgG2 gp41 | 0.068 | 0.322 | 1.000 |
| IgG3 gp41 | 0.024 | 0.728 | 1.000 |
| IgG4 gp41 | 0.249 | **0.000** | **0.004** |
| IgG C1 | 0.055 | 0.436 | 1.000 |
| IgG C5 | 0.054 | 0.433 | 1.000 |

**Supplementary Table 4.** Mutant pseudovirus panels for epitope mapping.

| Parent Virus | TRO11 | 25710-2.43 | BJOX002000.03.2 | Epitope Specificity |
| --- | --- | --- | --- | --- |
| Mutants | TRO.11.N279A | 25710-2.43.N279A |  | CD4bs |
|  | TRO.11.N280D | 25710-2.43.N280D |  |  |
|  | TRO.11.G458Y | 25710-2.43.G458Y |  |  |
|  | TRO.11.N276Q | 25710-2.43.N276Q |  |  |
|  | TRO.11.S365P |  |  |  |
|  | TRO.11.N295V |  | BJOX002000.03.2.L295N | 2G12 |
|  | TRO.11.N332A | 25710-2.43.N332A | BJOX002000.03.2.N332A | V3 Glycan |
|  | TRO.11.N160K | 25710-2.43.N160K | BJOX002000.03.2.N160K | V2 Glycan |
|  | TRO.11.N88A | 25710-2.43.N88A | BJOX002000.03.2.N88A | gp120-gp41 Interface |
|  | TRO.11.N625A | 25710-2.43.N625A | BJOX002000.03.2.N625A |  |
|  | TRO.11.N611A | 25710-2.43.N611A | BJOX002000.03.2.N611A |  |

**Supplemental Figure 5:**
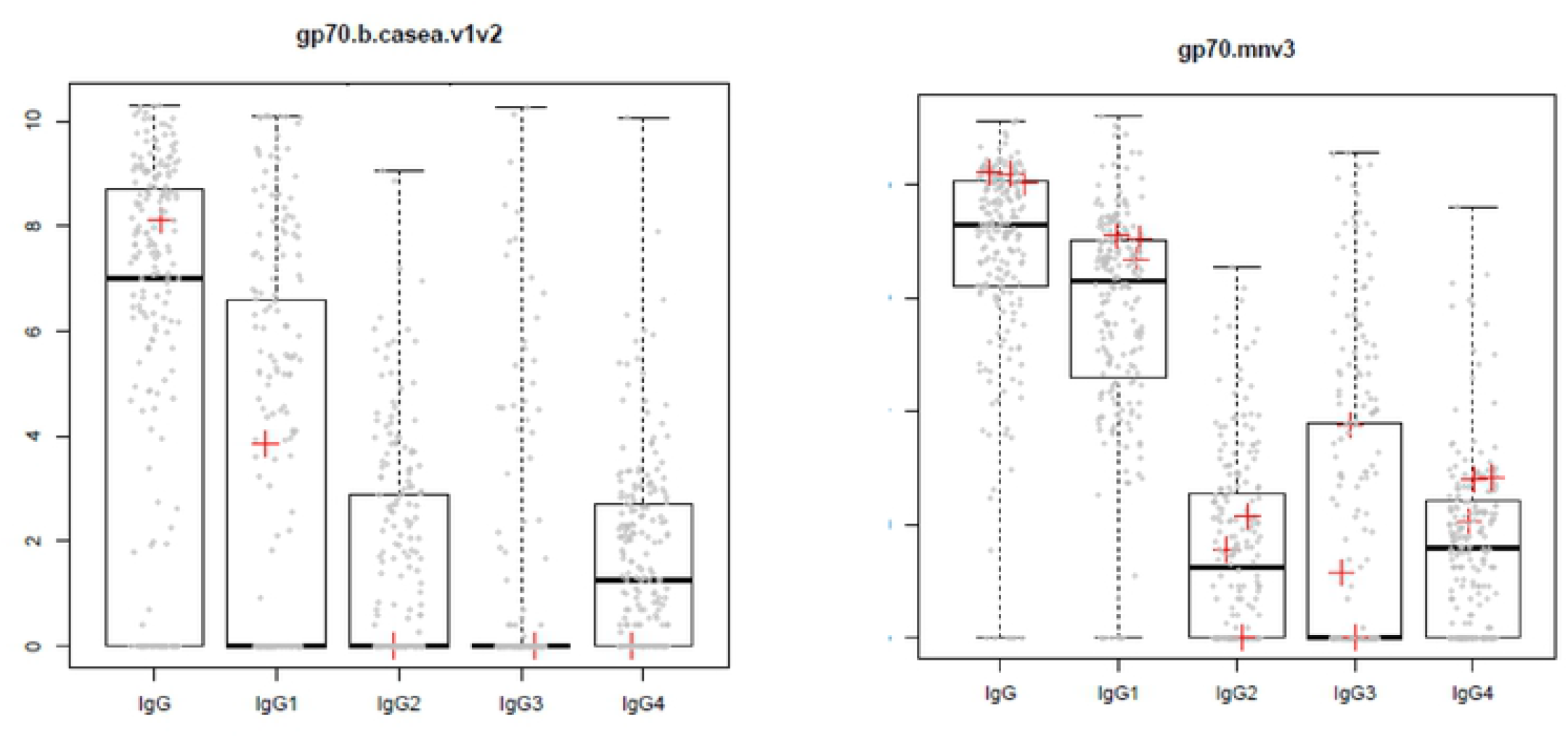
Association between binding and neutralization epitope specificity. Distributton of the V1V2-specific (A) and of V3-specific (B) binding responses in the pediatric cohort. The PTDS in whom neutralization specificity mapped to V2 glycan (PTD 1, panel A) or 10 V3 glycan (PTD 2,4 and 5, panel B) is indicated with the red cross. Overall. PTD ranks above the 50% percentile for V1V2 specific IgG and IgG1 but not for the other IgG subelasses. PTD 2,4,and 5 ranked above the 50% percentile for V3-specific IgG, IgG1 and IgG4.

